# Multiple changes underlie allelic divergence of *CUP2* between *Saccharomyces* species

**DOI:** 10.1101/728980

**Authors:** Xueying C. Li, Justin C. Fay

**Affiliations:** Molecular Genetics and Genomics Program, Washington University, St. Louis, MO; Department of Genetics, Washington University, St. Louis, MO; Center for Genome Sciences and System Biology, Washington University, St. Louis, MO; Department of Biology, University of Rochester, Rochester, NY

**Author notes:** Corresponding author: Xueying Li. European Molecular Biology Laboratory, Meyerhofstraße 1, 69117 Heidelberg, Germany (Xueying C. Li).

**Keywords:** *Saccharomyces*, *CUP2*, copper resistance, *cis*-regulatory evolution, chimeras

## Abstract

Under the model of micromutationism, phenotypic divergence between species is caused by accumulation of many small-effect changes. While mapping the causal changes to single nucleotide resolution could be difficult for diverged species, genetic dissection via chimeric constructs allows us to evaluate whether a large-effect gene is composed of many small-effect nucleotide changes. In a previously described non-complementation screen, we found allele difference of *CUP2*, a copper-binding transcription factor, underlie divergence in copper resistance between *Saccharomyces cerevisiae* and *S. uvarum*. Here, we tested whether the allele effect of *CUP2* was caused by multiple nucleotide changes. By analyzing chimeric constructs containing four separate regions in the *CUP2* gene, including its distal promoter, proximal promoter, DNA binding domain and transcriptional activation domain, we found that all four regions of the *S. cerevisiae* allele conferred copper resistance, with the proximal promoter showing the largest effect, and that both additive and epistatic effects are likely involved. These findings support a model of multiple changes underlying evolution and suggest an important role of both protein coding and *cis*-regulatory changes in evolution.

## Introduction

The genetic basis of evolutionary change may involve changes that range from large to small effect. Under the micromutational model, phenotypic divergence predominantly results from the accumulation of numerous small effect changes (Rockman 2011). However, mapping of quantitative traits has shown that large-effect changes often contribute to phenotypic variation (Orr and Coyne 1992; Bell 2009). Even so, these results may be inherently biased, both by a focus on dramatic phenotypic shifts, such as those that distinguish domesticated species from their wild relatives, and by the limited power of quantitative trait mapping to detect small effects and distinguish between regions with a single large-effect change or many small ones (Orr and Coyne 1992; Rockman 2011). Thus, evaluating the genetic basis of evolutionary change requires accounting for both the context and purview of the evidence.

In genetic studies, both the mapping method and samples size have a strong influence on the results. In contrast to many linkage mapping studies, which tend to find large-effect changes (Fay 2013), genome-wide association studies predominantly detect numerous small-effect associations, e.g. (Wood et al. 2014), and the number of associations depends on sample size (Visscher et al. 2012). Furthermore, evidence for the omnigenic model supports the view that every gene has some slight contribution to a trait (Boyle et al. 2017), and implies that the vast majority of causal variants are not realistically mappable. Knowing the limits of our ability to detect and identify small effect mutations is also relevant to answering questions about the genes, type of changes, and cellular mechanisms underlying phenotypic divergence (Rockman 2011; Boyle et al. 2017).

Limits on our ability to map phenotypic variation are not restricted to a simple tradeoff between effect size and sample size. Mapping interspecific differences often requires different approaches and yields different results compared to studies of intraspecific variation. A prominent limitation of mapping phenotypic differences between species is hybrid sterility and inviability. Consequently, many studies test candidate genes or map traits that differ between closely related, interfertile species. Based on a review of the literature, interspecific studies find fewer null alleles and more *cis*-regulatory alleles compared coding alleles (Stern and Orgogozo 2008). Another factor relevant to interspecific studies is that there is enough time for multiple changes to occur at a single locus. These loci are of interest both in regards to why they accumulate multiple changes, but also because they are more readily detected.

Repeated changes at a single locus, termed evolutionary hotspots, are common and relevant to understanding phenotypic divergence (Martin and Orgogozo 2013). Hotspots can be classified as interlineage, involving genes that are repeatedly used during evolution in different lineages, or intralineage, involving the accumulation of multiple changes in a gene along a single lineage (Martin and Orgogozo 2013). In the case of intralineage hotspots, multiple changes within a single gene can be explained by either the unique ability of a gene to affect a trait or pleiotropy, whereby many genes can influence a trait but relative few can do so without adverse effects on other traits (Stern and Orgogozo 2009). The constraints of pleiotropy are also thought to increase the preponderance of *cis*-regulatory changes in evolution (Carroll 2008). An example of one such hotspot is *shavenbaby*, which underlies divergence in trichomes between *Drosophila* species via multiple *cis*-regulatory changes (McGregor et al. 2007).

If phenotypic divergence between species results from the accumulation of numerous changes of small effect, they may be easiest to detect when they form hotspots. However, identifying hotspots between species is also a challenge. Species that are too close may not have enough time to accumulate multiple changes and species that are too distant may be reproductively isolated. Genetic analysis of species’ hybrids provides a means of balancing these limitations. Hybrids, even if infertile, are often viable for distantly related species. Hybrids have been leveraged for deletion mapping of incompatibilities between *Drosophila* species, e.g. (Coyne et al. 1998; Tang and Presgraves 2009), and for reciprocal hemizygosity analysis in *Saccharomyces* species (Weiss et al. 2018; Li et al. 2019). The reciprocal hemizygosity test compares two hybrids each with a different allele deleted, thereby testing for allelic differences while controlling for haploinsufficiency (Steinmetz et al. 2002). Of particular relevance, the test examines the combined effects of all regulatory or coding differences between the two species’ alleles.

In this study we test whether single or multiple changes underlie allelic divergence of *CUP2* between *Saccharomyces* species. Using a genome-wide non-complementation screen, we previously found that divergence of *CUP2* contributed to the evolution of copper resistance in *Saccharomyces* species (Li et al. 2019). *S. cerevisiae* can tolerate high concentration of copper sulfate, a stress associated with vineyard environments. Although the level of copper resistance is variable among *S. cerevisiae* strains (Fay et al. 2004; Kvitek et al. 2008; Strope et al. 2015), it’s relatives, *S. paradoxus* and *S. uvarum*, are usually copper sensitive (Kvitek et al. 2008; Warringer et al. 2011; Dashko et al. 2016). Through a non-complementation screen followed by a reciprocal hemizygosity test, we found that the *S. cerevisiae CUP2* allele confers higher copper resistance compared to the *S. uvarum* allele. *CUP2* encodes a copper-binding transcription factor and regulates Cup1p, a major copper-activated metallothionine in yeast (Buchman et al. 1989). Previous studies showed that *CUP2* is essential for *S. cerevisiae*’s copper resistance (Thiele 1988; Welch et al. 1989; Jin et al. 2008) and contributes to intraspecific variation in acetic acid (Meijnen et al. 2016) and copper resistance (Chang et al. 2013). Because the sequences of *S. cerevisiae* and *S. uvarum CUP2* are substantially diverged (71.1% identical) we dissected the effect of *CUP2* allele divergence using chimeric constructs between the two species. We found that divergence in copper-resistance is caused by multiple nucleotide changes distributed throughout the gene, but with *cis*-regulatory changes having a larger effect than coding changes.

## Materials and Methods

*S. cerevisiae* strains in the S288C background and *S. uvarum* strains in the CBS7001 background (Scannell et al. 2011) were used in this study. The *S. uvarum* genome sequence and annotations were from Scannell et al. (2011). *CUP2* was knocked out with *KanMX4* in *S. cerevisiae* (YJF173, *MAT***a** *ho-ura3-52*) and *S. uvarum* (YJF1450, *MAT***α** *hoΔ::NatMX*), respectively. Transformations in this study followed a standard lithium acetate procedure (Gietz et al. 1995), with the modification that room temperature and 37°C was used for incubation and heat shock of *S. uvarum*, respectively. Unless otherwise noted, *S. cerevisiae* was maintained at 30°C on YPD (1% yeast extract, 2% peptone and 2% dextrose) while *S. uvarum* and *S. cerevisiae* × *S. uvarum* hybrids were maintained at room temperature.

Chimeric constructs were generated by Gibson assembly (Gibson et al. 2009). Promoters were defined from the end of the upstream gene (*PMR1*) to the start codon of *CUP2*. Coding sequence (CDS) was defined from the start codon of *CUP2* to the stop codon, and our constructs also included the 3’ non-coding region (until the downstream gene). To further dissect the effects of the promoter and CDS, the promoter was split at nucleotide position −291 for *S. cerevisiae* and its homologous position at −283 for *S. uvarum*. The CDS was split at position +367 for both alleles, based on the previously defined DNA binding domain and transactivation domain (Buchman et al. 1989) (Fig. 1A). All positions are relative to the start codon of *CUP2.*

**Figure 1.**
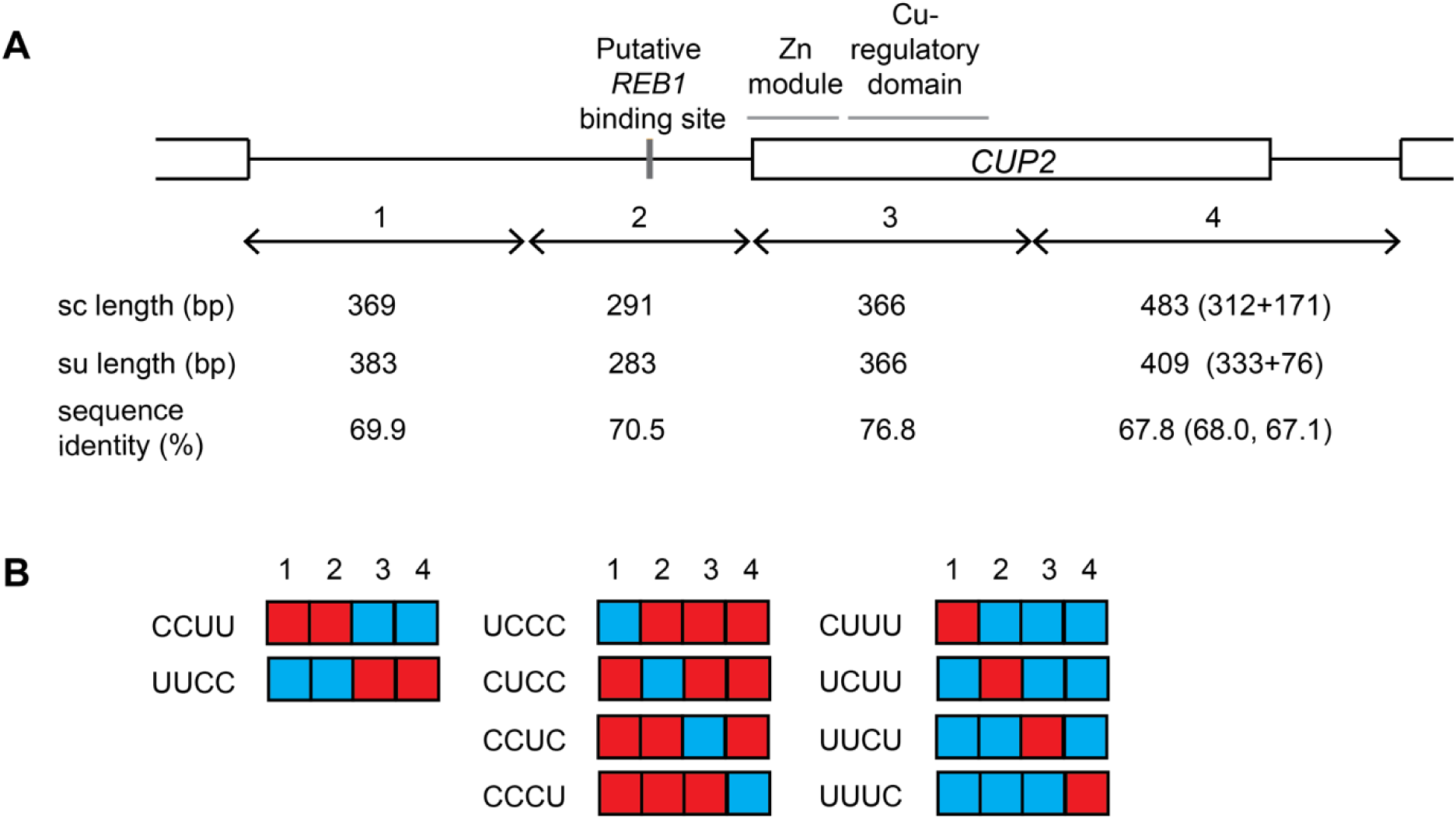
Design of *CUP2* chimeras. **A**. Diagram of *CUP2* gene, with black lines representing non-coding regions and boxes representing coding regions. The alleles were split into 4 regions (1-4). Region 2 contains a putative *REB1* binding site (De Boer and Hughes 2012) and region 3 contains the DNA binding domain (DBD) (Buchman et al. 1989), including a 40-residue zinc module (Turner et al. 1998) and a ∼60 residue copper regulatory domain (Graden et al. 1996). The diagram is drawn to scale of the *S. cerevisiae* allele, with the length of *S. cerevisiae* (sc) and *S. uvarum* (su) regions indicated below. Sequence identity is based on MUSCLE alignments, without counting gaps. Region 4 includes the 3’ half of the coding sequence and the 3’ intergenic sequence, of which the sequence length and identity was separately indicated in parentheses. **B**. *S. cerevisiae* (C, red) and *S. uvarum* (U, blue) segments were assembled into 10 chimeric constructs, including promoter-swaps (left), different *S. uvarum* regions inserted into the *S. cerevisiae* allele (middle), and different *S. cerevisiae* regions inserted into the *S. uvarum* allele (right).

Segments of *CUP2* were PCR-amplified from *S. cerevisiae* or *S. uvarum* genomic DNA with Q5 polymerase (New England Biolabs). Promoter and CDS segments from different species were Gibson-assembled into pRS306 to generate promoter-swaps. Full-length *S. cerevisiae* and *S. uvarum CUP2* alleles were assembled in parallel for controls. An *S. cerevisiae* allele from a copper sensitive oak tree strain was included for comparison, and was amplified from genomic DNA of YJF153 (*MAT***a** *hoΔ::dsdAMX*), a YPS163 derivative. To split the promoter or CDS, the segments of interest were assembled into pRS306-derived plasmids pXL07 or pXL05, which respectively carry the full-length *S. cerevisiae* or *S. uvarum* allele. All constructs were Sanger-sequenced; one of the chimeras (CCUC) carried a deletion of a single adenine nucleotide in a stretch of 14 As in the *S. cerevisiae* promoter, but it did not seem to cause deleterious effects in the phenotypic assays.

The plasmids were linearized with BstBI (*CUP2* constructs) or StuI (vector control) and integrated into the *ura3* locus of an *S. cerevisiae CUP2* knockout strain YJF2872 (*MAT***a** *ho-ura3-52 cup2Δ::KanMX4*). The integrated strains were backcrossed to an *S. cerevisiae* strain YJF175 (*MAT***α** *ho-ura3-52*) and sporulated to remove any second-site mutations. The resulting haploid *S. cerevisiae* strains carrying the *CUP2* deletion and chimeric constructs were then crossed to an *S. uvarum CUP2* knockout YJF2917 (*MAT***α** *hoΔ::NatMX cup2Δ::KanMX4*). The final interspecific hybrid was null for both *S. cerevisiae* and *S. uvarum* alleles at their endogenous loci and carried chimeric or full-length constructs at the *ura3* locus. The hybrids were genotyped by PCR (Li et al. 2019) and found to carry *S. cerevisiae* mitochondrial DNA.

Growth curves in copper-supplemented media were recorded by a BioTek microplate reader. Three biological replicates were used for each strain. Overnight cultures were diluted 1:100 into 200 ul complete media (CM, 0.3% yeast nitrogen base with amino acids, 0.5% ammonium sulfate, 2% dextrose) supplemented with 0, 0.2 or 0.5mM copper sulfate in a 96-well plate. The plate was incubated at room temperature (25-26°C), with the optical density (OD) at 600 nm taken every 10min for 40h. The plate was shaken for 20s before each OD reading. To quantify growth differences, area under the curve (AUC) was measured as the integral of the spline fit of growth curves using the grofit package (Kahm et al. 2010) in R. Copper resistance was represented by normalized AUC (nAUC), the AUC of copper treatments divided by the mean AUC of the same strain in CM without copper.

Linear models were used to analyze the effects of each region. Data from the oak allele and the vector control were excluded in the models. The sum of nAUC across the two concentrations (snAUC) was used to represent copper resistance of each strain. The data were fit to two models: 1) *snAUC ∼ R1 + R2 + R3 + R4*, to analyze the additive effects of region 1 to 4 (*R1* to *R4*); 2) *snAUC ∼* (*R1 + R2 + R3 + R4*) ^2, to analyze both additive and epistatic effects. *R1* to *R4* were categorical variables (C or U representing *cerevisiae* and *uvarum* alleles, respectively). P-values were extracted from the models and were adjusted by false discovery rate (Benjamini and Hochberg method) to correct for multiple comparisons. All data and reagents used in this study are available upon request.

## Results

The *S. cerevisiae* allele of *CUP2* confers higher copper resistance than the *S. uvarum* allele (Li et al. 2019). The two alleles share 71.1% sequence identity, with hundreds of nucleotide substitutions across the coding and non-coding regions. To test whether the allele differences in copper resistance are caused by multiple nucleotide changes and whether they occur in coding or *cis*-regulatory regions, we generated chimeric constructs between *S. cerevisiae* and *S. uvarum CUP2* alleles (Fig. 1) and integrated them into the *ura3* locus in *S. cerevisiae*. Copper resistance was measured in a hybrid of *S. cerevisiae* and *S. uvarum*, in which the endogenous *CUP2* alleles were knocked out. The hybrid background was used in accordance with the previously conducted reciprocal hemizygosity test (Li et al. 2019), but the effects of chimeras were the same in *S. cerevisiae* (Fig. S1).

All four of the regions showed a significant effect on copper resistance using an additive model (Table 1). Across two different concentrations of copper, the resistance of chimeras generally increased with the number of *S. cerevisiae* segments in the constructs (Fig. 2). Relative to the *S. uvarum* allele, substituting in the *S. cerevisiae* promoter conferred higher resistance than substituting the *S. cerevisiae* CDS (grey). The chimeras that split the promoter or CDS regions further mapped the largest effect to the proximal half of the *S. cerevisiae* promoter (the UCUU construct), while the other three *S. cerevisiae* regions tested also conferred low-to-moderate levels of resistance when inserted into the *S. uvarum* allele (light blue, left panel), suggesting that multiple nucleotide changes underlie the allele effect of *CUP2.* While the combination of any three *S. cerevisiae* segments was sufficient to confer resistance to the 0.2mM copper treatment (orange), these chimeras showed various levels of sensitivity to 0.5mM, also consistent with a model of multiple changes.

**Table 1.** Additive and epistatic effects of *S. cerevisiae CUP2* regions on copper resistance.

| Region<sup>#</sup> | Additive model |  | Epistatic model |  |
| --- | --- | --- | --- | --- |
|  | Effect size | P-value<sup>†</sup> | Effect size | P-value<sup>†</sup> |
| (Intercept) | 0.138 | 0.0841 | 0.197 | 0.000445 |
| 1 | 0.479 | 3.11E-06 | 0.314 | 9.37E-05 |
| 2 | 0.515 | 1.33E-06 | 0.801 | 1.29E-11 |
| 3 | 0.274 | 0.00267 | 0.333 | 5.59E-05 |
| 4 | 0.527 | 1.33E-06 | 0.211 | 0.00369 |
| 1*2 |  |  | -0.339 | 0.000292 |
| 1*3 |  |  | 0.0754 | 0.370 |
| 1*4 |  |  | 0.594 | 1.47E-07 |
| 2*3 |  |  | -0.232 | 0.00679 |
| 2*4 |  |  | NA | NA |
| 3*4 |  |  | 0.0381 | 0.622 |

<sup>#</sup> Regions were defined as in Fig. 1A. The asterisks indicate interactions.
<sup>†</sup> P-values were adjusted by the false discovery rate (Benjamini and Hochberg method).

**Figure 2.**
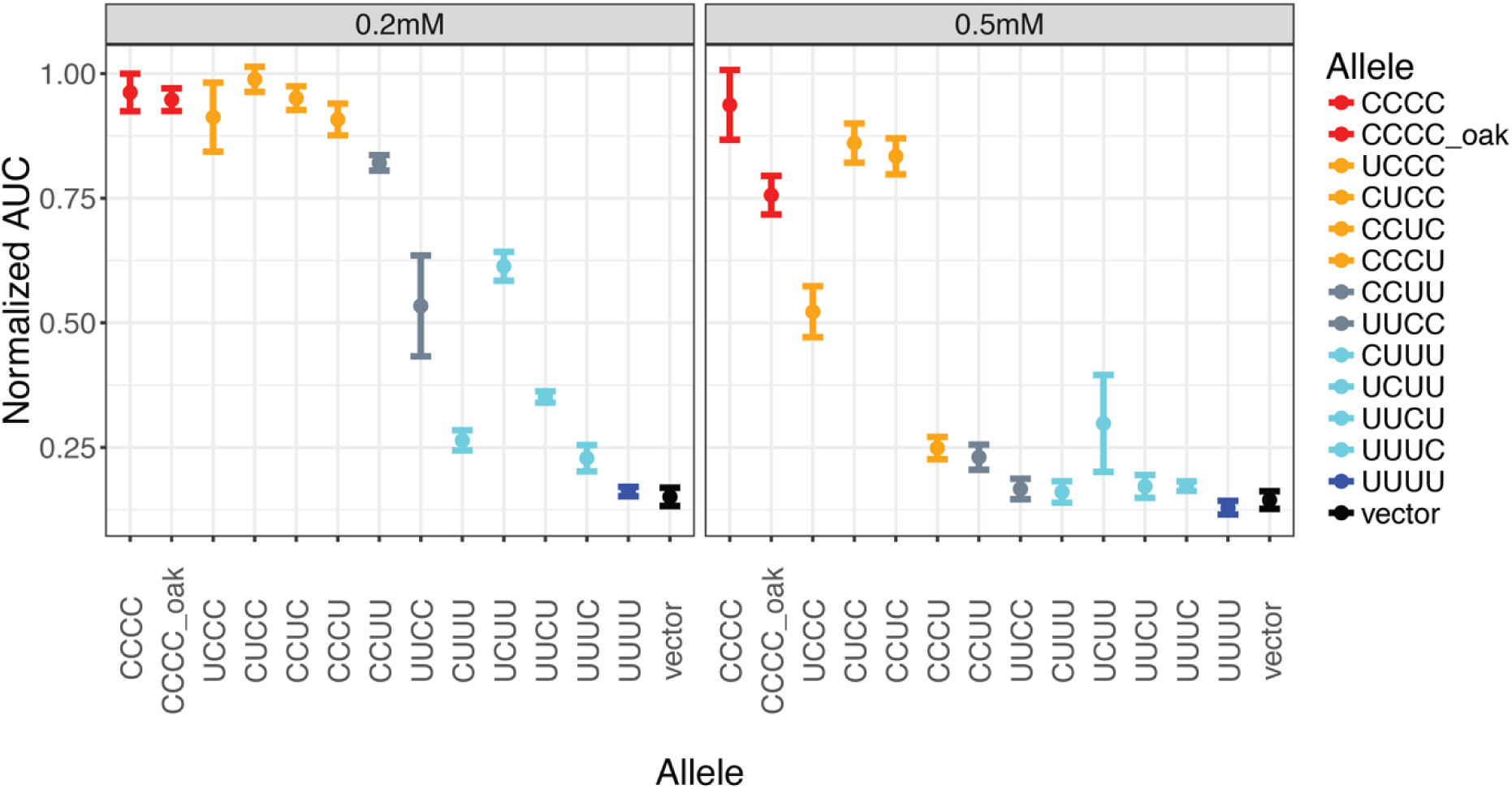
Copper resistance of chimeric constructs. *S. cerevisiae* × *S. uvarum* hybrids carrying the chimeric constructs were grown in labeled copper concentrations and their resistance was measured by area under curve (AUC) of OD_600_ growth curves, normalized to their growth in complete media. Points represent the mean of three biological replicates and error bars represent 95% confidence interval. The colors are based on the number of *S. cerevisiae* segments in the chimeras (red = 4, orange = 3, grey = 2, light blue = 1, blue or black = 0).

Using a linear model we also tested whether there are epistatic interactions between the regions (Table 1). We found that the model accounting for epistatic effects explained the data better than the model with only additive effects (0.974 vs. 0.839 for adjusted R-squared, p=1.94E-10 in ANOVA). In the epistatic model, all four *S. cerevisiae* regions retained significant effects on copper resistance, with region 2 showing the largest effect. Positive epistasis was detected between region 1 and 4. At high copper concentration, substitution of *S. cerevisiae* region 1 or 4 into the *S. uvarum* background had little effect (Fig. 2, right panel, CUUU and UUUC compared to UUUU), but showed much larger effects when the other region was also present (CCUU to CCUC and UUCC to CUCC). Regions 1-2 and 2-3 showed modest negative interactions. These findings suggest that both changes with additive and epistatic effects contributed to the divergence of *CUP2* alleles.

We also included a full-length *CUP2* allele from a copper-sensitive *S. cerevisiae* oak isolate for comparison. The oak allele has 12 nucleotide differences from the S288C allele used in the chimeras. While the oak allele showed similar levels of resistance as the S288C allele at 0.2 mM copper, it was more sensitive than the S288C allele at 0.5 mM. This suggests that a portion of the divergence between the *S. cerevisiae* S288C allele and *S. uvarum* may be caused by recent changes (polymorphism). However, of the 572 differences between the S288C and *S. uvarum* allele (out of a 1586 bp alignment, including gaps), only 4 of these can be explained by polymorphism between the two *S. cerevisiae* strains and only 57 of these are polymorphic in other *S. cerevisiae* strains (Peter et al. 2018).

## Discussion

Evolution can occur through accumulation of many small-effect changes, but mapping small-effect changes can be technically challenging (Orr 2001; Rockman 2011). In the present study, we tested whether a relatively large effect on copper resistance caused by *CUP2* allele divergence is a consequence of multiple nucleotide changes. By splitting the *CUP2* gene into four regions and measuring their effects via chimeric constructs, we found that the *CUP2* allele difference was caused by accumulation of multiple small-to-medium effect changes, with the proximal promoter region showing the largest effect.

### Multiple changes with small effects

Our findings support the micromutationism view that evolution involves many small-effect changes. All four regions tested conferred copper resistance with various effect sizes, suggesting that the copper-resistant nucleotide substitutions are distributed throughout the *CUP2* gene. The largest effect was mapped to the proximal promoter. The promoter effect was unlikely to be caused by changes in transcription factor binding sites: there is only one putative *REB1* binding site in the *CUP2* promoter (YetFasCo database, (De Boer and Hughes 2012), Fig. 1A), and it is conserved across the *Saccharomyces* species. The large effect of the *CUP2* promoter supports the previously suggested prominent role of *cis*-regulatory changes in long-term evolution (Stern and Orgogozo 2008). While *cis*-regulatory changes were often found to underlie morphological evolution, the example of *CUP2* along with several prior studies demonstrated that they are also important to physiological traits in yeast (Gerke et al. 2009; Engle and Fay 2012; Roop et al. 2016).

Cup2p consists of an N-terminal DNA binding domain (region 3) and a C-terminal transcriptional activation domain (region 4) (Buchman et al. 1989), with the former being more conserved (Fig. 1A). We found that the DNA binding domain of *S. cerevisiae* conferred moderate copper resistance when inserted into the *S. uvarum* allele. The gain of copper resistance could be due to changes in binding affinity to the *CUP1* promoter, the major target of Cup2p. The N-terminal of Cup2p is suggested to bind DNA via a zinc module and a copper-regulatory domain (Graden et al. 1996) (Fig. 1A), both of which contain amino acid differences between the two species. Further dissection of this region would help understand the molecular mechanism of *CUP2*-mediated copper resistance. However, these dissections are expected to become increasingly difficult under the micromutational model.

While all four regions showed different levels of additive effects, the context-dependent effect sizes of individual regions suggest epistasis. The *S. cerevisiae* region 1 and 4 showed small effects when inserted into the *S. uvarum* allele (Fig. 2, CUUU and UUUC constructs) but large effects when replaced by the *S. uvarum* regions (Fig. 2, UCCC and CCCU). It is possible that these two regions of *S. cerevisiae* contain large-effect copper-resistant changes that depend on the presence of other *S. cerevisiae* regions. Alternatively, the *S. cerevisiae* region 1 and 4 may only contain small-effect changes, and the sensitivity of the UCCC and CCCU constructs was caused by deleterious effects of the *S. uvarum* regions. Our data could not distinguish these two possibilities, although the linear model suggested that synergistic epistasis between the *S. cerevisiae* region 1 and 4 could be the best explanation (Table 1).

### Evolution of copper resistance

The evolutionary history of *CUP2* provides some insight into the evolution of copper resistance. The *CUP2* coding sequences do not exhibit signatures of positive selection according to site-specific dN/dS models (Scannell et al. 2011) or McDonald-Kreitman tests (Doniger et al. 2008). However, the coding sequences do show significant heterogeneity in the dN/dS ratio across *Saccharomyces* lineages (p=0.00523 compared to a model of fixed rates), indicating variation in selection pressure across lineages, with the *S. cerevisiae* lineage showing the highest ratio (0.562) (Scannell et al. 2011). The gain of copper resistance of *S. cerevisiae* has been associated with its adaptation to vineyard environments, where copper has been used as a fungicide (Mortimer 2000). While this trait is variable within *S. cerevisiae*, suggesting recent adaptation, most tested strains of *S. paradoxus* and *S. uvarum* are sensitive (Kvitek et al. 2008; Warringer et al. 2011; Dashko et al. 2016). Therefore, *S. cerevisiae* might have acquired copper-resistant changes prior to adaptation of wine strains to the vineyard. This view is supported by the observation that the *S. cerevisiae* oak allele, which is from one of the most copper sensitive *S. cerevisiae* strains (Fay et al. 2004), showed much higher copper resistance than the *S. uvarum* allele of *CUP2*. While variation in copper resistance within *S. cerevisiae* strains is largely attributed to copy number variation of *CUP1* and *CUP2* (Fogel and Welch 1982; Chang et al. 2013), the interspecific divergence may have a more complex genetic architecture. We showed that multiple changes in *CUP2* contribute to copper resistance in the present study, but the sum of their effects did not account for the total difference between *S. cerevisiae* and *S. uvarum* (Li et al. 2019). Fully elucidation of the genetic basis of copper resistance would require further genetic analysis between *Saccharomyces* species.

## Acknowledgments

We thank members of Fay lab for comments and experimental assistance. This work was supported by a National Institutes of Health grant (GM080669) to J.C.F.

---

**Figure S1.**
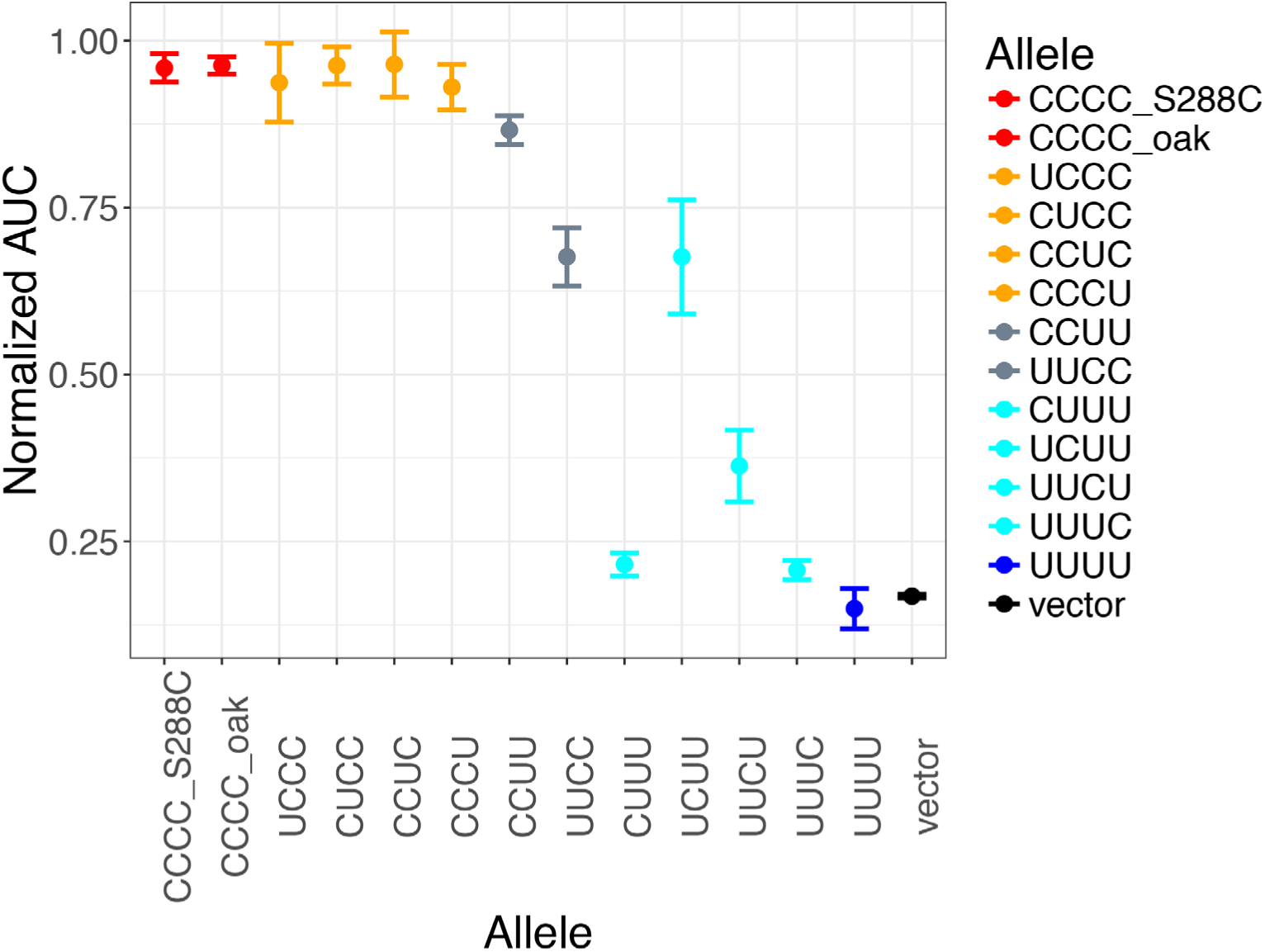
Copper resistance of chimeric constructs in *S. cerevisiae.* Copper resistance of the chimeras was examined in an *S. cerevisiae CUP2* knockout strain in 0.5 mM copper sulfate. Resistance was measured by normalized area under the curve (AUC), with points representing the mean of three biological replicates and error bars representing 95% confidence interval. The colors are based on the number of *S. cerevisiae* segments in the chimeras (red = 4, orange = 3, grey = 2, light blue = 1, blue or black = 0).

